# Engineered Proteins Mimic ssRNA Phage to Disrupt Type IV Pili

**DOI:** 10.64898/2025.12.14.694263

**Authors:** Addison Frese, Zhi Zhao, Lanying Zeng

## Abstract

Antimicrobial resistance is a growing global health crisis, with ESKAPE pathogens, such as *Acinetobacter* species, contributing significantly to hospital-acquired infections. These bacteria employ various virulence factors, including extracellular type IV pili (T4P), which serve as essential appendages for DNA uptake, biofilm formation, and resistance to antibiotics. Single-stranded RNA (ssRNA) bacteriophages (phages) exploit these retractile pili as entry receptors, but the exact mechanism of viral RNA delivery following pilus binding and retraction remains poorly understood. In this study, we investigated the entry mechanism of the ssRNA phage AP205 by identifying the specific phage components necessary for T4P detachment, a key step in viral genome entry. Our data reveals the phage’s maturation protein (Mat) is sufficient to induce pilus detachment on its own, and the overall virion structure further enhances the efficiency of this process. These findings provide mechanistic insights into how ssRNA phages exploit bacterial T4P as part of their infection pathway and suggest a conserved strategy for host entry.

**Importance:** *Acinetobacter* species are widespread environmental bacteria and members of the ESKAPE group of pathogens, often associated with hospital-acquired infections amongst immunocompromised patients. A major reason for the increase in antibiotic resistance in *Acinetobacter* species is its use of T4P to evade antibiotic treatment. With antibiotic resistance becoming a global health crisis, targeting these virulence factors represents a promising approach to combat infections. Since T4P are critical for *Acinetobacter* virulence, understanding mechanisms that can disrupt them is highly important. ssRNA phages provide a natural model for this, as they can target T4P to initiate infection. Recent research has shown that ssRNA phages can detach pili during their infection process. However, the precise mechanism behind this phenomenon remains unknown. Our research addresses part of this knowledge gap by identifying the key components required for this pilus detachment. This work will enhance the current understanding of the virulent T4P in *Acinetobacter* species, which could ultimately lead to broader medical advancements.

## Introduction

The global healthcare system faces a formidable challenge from the escalating prevalence of hospital-acquired infections, primarily driven by the relentless emergence and spread of antimicrobial-resistant (AMR) bacterial strains. Among these, *Acinetobacter* species (spp.) have emerged as a significant contributor to AMR bacterial human diseases. This is underscored by a staggering 78% increase in nosocomial infections attributed to *Acinetobacter* spp. between 2019 and 2020^1^. A particularly concerning member of this genus, *Acinetobacter baumannii*, has exhibited a marked and alarming increase in resistance to multiple classes of antibiotics. Over several years, the proportion of *A. baumannii* isolates resistant to at least three antibiotic classes has surged dramatically from 4% to 55%^2^. The remarkable adaptability of *Acinetobacter spp*., gram-negative bacteria capable of surviving in diverse environments such as soil and water, has enabled their insidious rise to become an urgent threat to patient health, especially within hospital-like settings where vulnerable populations are concentrated^3,4^.

A key factor in the virulence and adaptability of *Acinetobacter* spp. is their utilization of extracellular appendages known as type IV pili (T4P). T4P are hair-like, filamentous structures that exhibit dynamic cycles of extension and retraction that facilitate their involvement in a wide array of essential cellular processes, including motility, biofilm formation, adhesion to host cells, and the uptake of genetic materials, which contribute to the spread of antibiotic resistance genes^5,6^. T4P are commonly composed of thousands of copies of PilA, the major pilin subunit, which form helical polymers with a few minor pilin subunits that cap the pilus^7–9^.

Single-stranded RNA bacteriophages (ssRNA phages) infect bacteria by exploiting bacterial pili. ssRNA phages have small genomes of ∼4000 nucleotides, containing a single positive-sense viral RNA (vRNA) strand, which encodes four essential proteins: maturation protein (Mat), coat protein (Coat), replicase (Rep), and single-gene lysis (Sgl) protein^11^. The virion consists of 178-180 copies of Coat, which encapsulate the vRNA. Typically, the vRNA is bound to Mat, which protrudes from the viral capsid, serving as the initial point of contact with a host pilus^12^. A key model for studying these interactions in *Acinetobacter* species is the ssRNA phage AP205, as it uniquely possesses two identical Mat proteins outside of the capsid^13,14^. To isolate the function of its pilus-binding domain, previous research has engineered a truncated construct of the Mat protein, called Mat_200_, and fused it with superfolder green fluorescent protein, generating Mat_200_-sfGFP^10^. Mat_200_ is composed of the first 200 N-terminal amino acids of the native AP205 Mat protein, which was shown to be sufficient to mediate binding to *Acinetobacter higginsii* (*A. higginsii*, NCBI:txid70347) T4P^10^. For ssRNA phages, infection begins once the phage binds to a pilus via Mat^11,15^. The canonical model of ssRNA phage infection requires pilus retraction to bring the phage to the cell surface, delivering the vRNA into the bacterial cell through a mechanism yet to be fully understood^15,16^. Intriguingly, it was recently discovered that the *E. coli* F-pili and *P. aeruginosa* T4P are detached from the cell surface once the pilus-bound ssRNA phages MS2/Qβ and PP7 are brought in proximity to the basal body of the cell, respectively^12,13^.

In this study, we built upon our pilus detachment model to define the phage components that are required for this event. We first expanded the model’s scope by showing the ssRNA phage AP205 also detaches *A. higginsii* T4P, pointing to a universally conserved mechanism. We then identified the Mat protein as the minimal component sufficient for detachment, which was tested in two different phage-host systems: Mat_200_ with *A. higginsii* T4P and the Qβ Mat protein with *E. coli* F-pili. Critically, we discovered that the Mat protein alone is less efficient than the whole phage, and that efficiency could be fully restored in *A. higginsii* by attaching Mat_200_ to a nanoparticle mimicking the size and rigidity of AP205 coat proteins. Our findings thus reveal that efficient pilus detachment is a two-part mechanism, dependent not only on the specific binding of the Mat protein but also on the physical leverage provided by the intact ssRNA phage capsid.

## Results

### AP205 causes the detachment of *A. higginsii* T4P

Previous work found that ssRNA phages MS2/Qβ and PP7 detach *E. coli* F-pilus and *P. aeruginosa* T4P during the entry process, respectively^12,13^. To determine if this is a universal phenomenon for ssRNA phages and retractile pili, we investigated the effect of the ssRNA phage AP205 on the T4P of its host, *A. higginsii*^14,15^. To visualize and quantify the T4P of *A. higginsii*, we utilized a previously constructed fluorescent protein fusion Mat_200_-sfGFP^10^. As expected, Mat_200_-sfGFP specifically binds to the T4P, illuminating them as long filaments extending from the cells (**Fig. 1A**). In the absence of *A. higginsii* T4P, Mat_200_-sfGFP signal appears as diffuse, homogeneous fluorescence (**Fig. S1**), confirming that Mat_200_-sfGFP specifically interacts with T4P. Remarkably, AP205 infection led to a marked increase in the number of free T4P in the media compared to the buffer-only control. The frequency of detached pili increased over time, reaching a plateau of ∼0.3 detached pili per cell after 20 min post-infection at a multiplicity of infection (MOI) of 5 (**Fig. 1B**). The buffer treated cells had only ∼0.03 detached pili per cell which did not change over time. This observation aligns with previous findings for the ssRNA phage PP7, which infects *P. aeruginosa*^12,13^.

**Figure 1:**
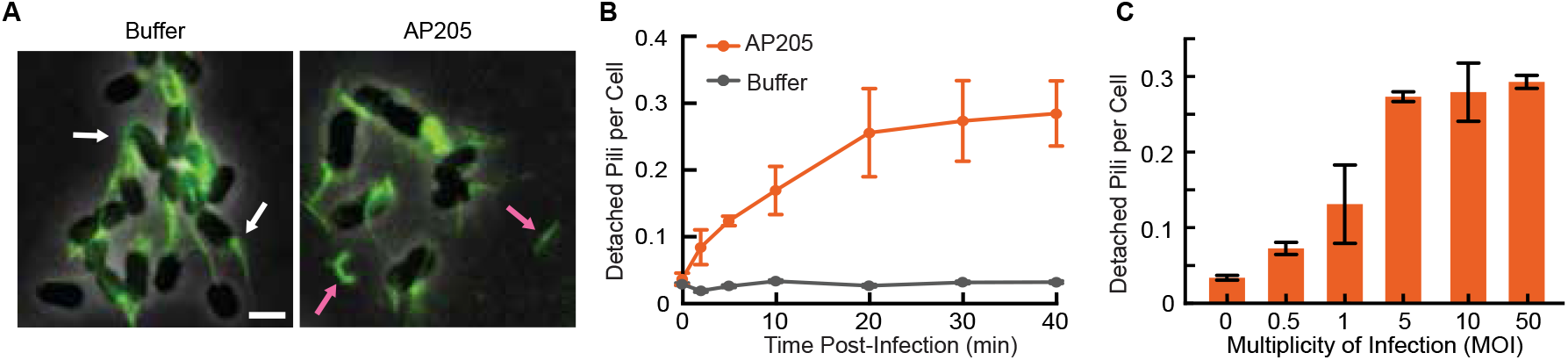
AP205 detaches *A. higginsii* T4P upon infection. (A) Representative images illustrating cell-associated pili (indicated by white arrows) and detached T4P from *A. higginsii* cells (indicated by pink arrows). (B) The number of detached pili per cell increases over the initial 20 min, reaching a plateau of ∼0.3 pili per cell upon AP205 infection, while the detachment remains ∼0.03 pili per cell for the buffer-treated sample. The total number of detached pili per cell at 0, 2, 5, 10, 20, 30, and 40 min were for AP205, 62/2128, 70/904, 105/855, 359/2159, 495/2132, 237/949, and 162/627, respectively; and for buffer, 51/1944, 62/3366, 50/1837, 178/5022, 123/4897, 104/2514, and 59/1789, respectively. Detached pili per cell were quantified by dividing the total number of detached pili by the total number of cells. Experiments were performed in at least triplicate. (C) The number of detached pili per cell increases with MOI, plateauing at ∼0.3 pili per cell at an MOI of 5 or greater. The mean number of detached pili per cell for MOI of 0, 0.5, 1, 5, 10, and 50 was for AP205, 36/493, 42/378, 85/312, 73/264, and 98/334, respectively; and for buffer, 25/711. Experiments were performed in at least triplicate. Scale bar, 2 μm.

We next investigated how phage MOI affects the frequency of T4P detachment. Cells were infected with AP205 at MOIs of 0, 0.5, 1, 5, 10, and 50, and the number of detached pili per cell was quantified at 30 min post-infection, at which time the pilus detachment had plateaued. Notably, the number of detached pili per cell increased with MOI, reaching a plateau of ∼0.3 detached pili per cell at an MOI of 5 or greater (**Fig. 1C**). This suggests that a critical number of phage particles is required to trigger maximal pilus detachment, further supporting that this phenomenon is a direct result of the phage infection process. These results suggest that T4P detachment is likely a universally conserved phenomenon that plays a key role in the infection process across different host species.

### Mat_200_ alone is sufficient to detach T4P

Previous research using computational modeling suggested that pilus detachment was initiated by the pilus bending upon phage Mat entry into the outer membrane complex of the T4P machinery powered by pilus retraction^12^. Here, with the result of AP205 phage readily detaching the T4P of *A. higginsii* (**Fig. 1**), as other ssRNA phages do, we asked if Mat_200_ alone is sufficient to induce pilus detachment, which can be tested experimentally. To test this, we used the previously generated Mat_200_ from the Mat protein of AP205^10^ (**Fig. 2A** and **B**). Indeed, Mat_200_ at a concentration of 20 μM caused an increase in the number of free pili in the media (**Fig. 2C**). The number of detached pili per cell plateaued at ∼0.1 after 20 min of treatment, significantly higher than the buffer-treated sample, indicating that Mat_200_ is sufficient to induce T4P detachment (**Fig. 2D**). The Mat_200_ protein fusion includes a His-SUMO tag on the N-terminus, which aids in the purification and stability of the protein. To exclude the possibility that this tag influences Mat_200_ activity, we tested a purified SUMO tag as a control. The SUMO tag alone did not induce T4P detachment, confirming that the observed activity is solely attributable to the Mat_200_ protein (**Fig. S2**).

**Figure 2:**
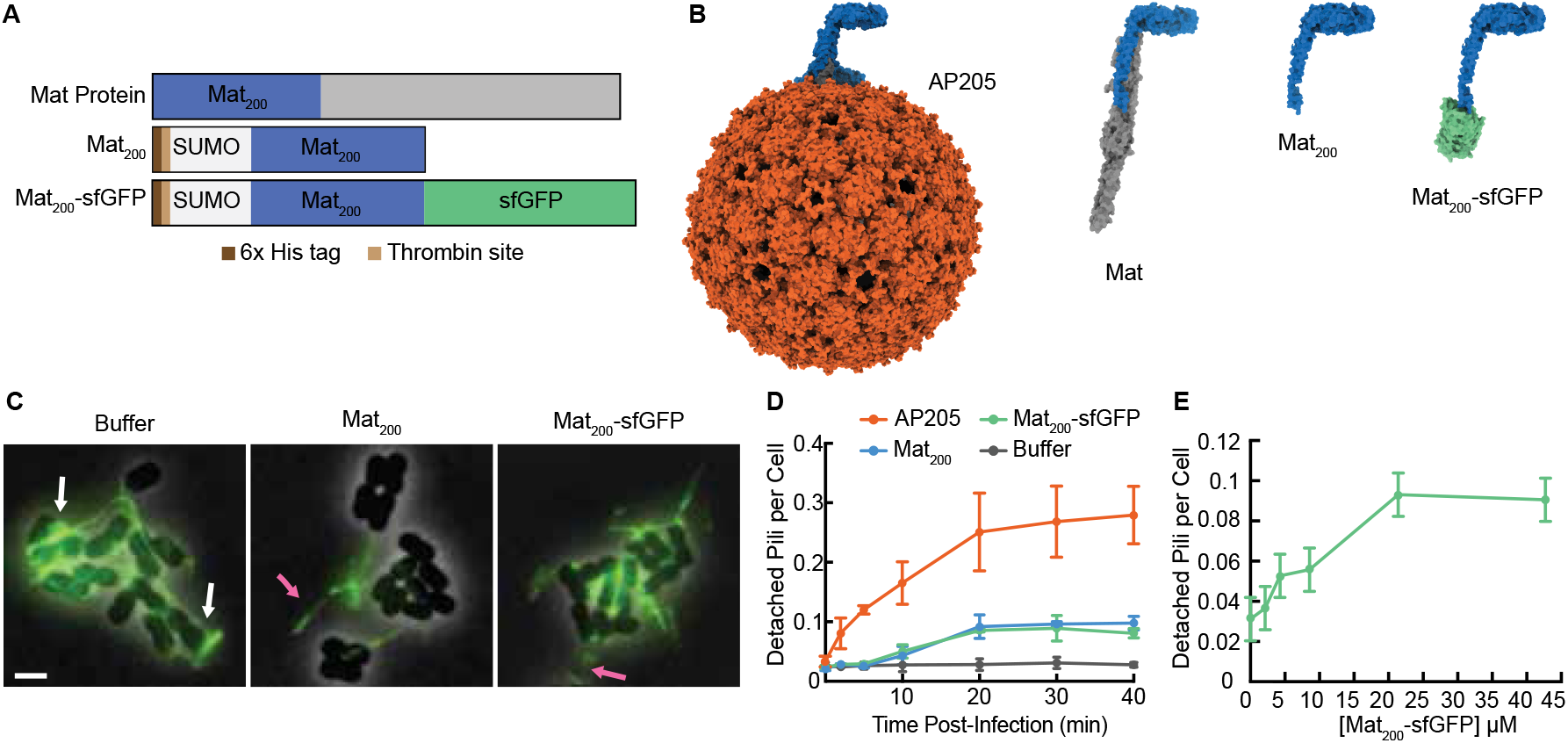
Mat_200_ and Mat_200_-sfGFP sufficiently detach T4P. (A) Comparison of the maturation protein (Mat), its truncated derivative Mat_200_ and Mat_200_-sfGFP. (B) Structural comparison of AP205, Mat, Mat_200_, and Mat_200_-sfGFP based upon previously published PDB entries for AP205 (8TOC), Mat_200_ (8TVA), and sfGFP (2B3P). The coat protein of AP205 is shown in orange. The Mat_200_ domain is consistently colored blue in all structures. For the full-length Mat protein (left), the remainder of the protein is shown in grey. In the Mat_200_-sfGFP construct (right), the sfGFP is depicted in green. Structures are shown to scale. (C) Representative micrographs demonstrating T4P detachment from *A. higginsii* cells. T4P were visualized by labeling with Mat_200_-sfGFP. White arrows indicate intact pili, whereas pink arrows highlight detached pili. (D) Quantification of detached pili per cell via various treatments over a time course of 0, 2, 5, 10, 20, 30, and 40 min post-treatment. Detached pili per cell increases over time for Mat_200_ and Mat_200_-sfGFP treated cells, plateauing at ∼0.1 detached pili per cell 20 min post-treatment. The total number of detached pili per cell for each treatment time was for Mat_200_, 31/1474, 42/1459, 36/1299, 93/2200, 144/1632, 243/2542, and 185/1854, respectively; for Mat_200_-sfGFP, 90/3824, 53/1872, 58/2009, 185/3473, 434/5408, 458/4672, and 376/4464, respectively; for buffer, 37/1479, 49/1775, 37/1362, 38/1852, 45/1871, 48/2023, and 37/1689, respectively. The data for AP205 treated cells was taken from **Fig. 1B**. Detached pili per cell was calculated as the total number of detached pili divided by the total number of cells in each experiment. Each experiment was performed in at least triplicate. (E) This graph illustrates the number of detached pili per cell after 30 min of treatment by various concentrations of Mat_200_-sfGFP. The total number of detached pili per cell for Mat_200_-sfGFP concentrations (∼0, 2, 4, 8, 21, and 42 μM) was 58/1631, 151/3472, 77/1711, 158/2936, 90/858, and 132/1384, respectively. Each concentration and dilution was tested in at least triplicate. Scale bar, 2 μm.

To explore the properties of Mat_200_ further, we examined whether the addition of a cargo protein would interfere with its function. We tested the pilus detachment by Mat_200_-sfGFP, the protein fusion used to visualize the pili previously^10^ in **Fig. 1**. Mat_200_-sfGFP at 20 μM also caused a significant increase in the number of detached pili, reaching a plateau of ∼0.1 detached pili per cell after 20 min, similar to the results with Mat_200_ alone (**Fig. 2D**). This observation demonstrates that the sfGFP cargo did not interfere with the ability of Mat_200_ to recognize and bind to the T4P and that it retained a similar efficiency of pilus detachment. Note that pilus detachment by Mat_200_-sfGFP does not occur at 2 min, at which time we have used to illuminate the pili. We next investigated the effect of Mat_200_-sfGFP concentration on pilus detachment. The number of detached pili per cell increased with the concentration of Mat_200_-sfGFP, reaching a plateau of ∼0.1 detached pili per cell at a concentration of 20 μM (**Fig. 2E**). This concentration-dependent behavior further confirms that the observed pilus detachment is a direct result of the binding of Mat_200_-sfGFP to the pili.

### Qβ Mat causes the detachment of *E. coli* F-pilus

To determine if the Mat protein’s pilus-detaching ability is a broad-spectrum phenomenon, we investigated another ssRNA phage system: the Qβ phage infecting its host, *E. coli*, via the conjugative F-pilus. Specifically, we examined the F-pilus detachment by a soluble Qβ maturation protein fused to a maltose-binding protein (MBP), referred to as MBP-Mat_Qβ_ ^16^. To visualize F-pili, we used previously established fluorescent MS2-GFP phages and treated *E. coli* HfrH cells with MBP-Mat_Qβ_ (**Fig. 3A**)^13^. Compared to the buffer-only control, cells treated with 1 μM MBP-Mat_Qβ_ showed a significant increase in the number of detached pili in the media (**Fig. 3A**). The percentage of detached pili over total pili reached a plateau of ∼17% after 10 min of treatment, which is lower than the ∼26% plateau observed in cultures infected with intact Qβ phages (**Fig. 3B**). The buffer-treated cells had only ∼5% detached pili, which did not increase with time. The detachment of buffer or Qβ-treated cells is consistent with that of previously published data^13^. These results indicated that despite less efficient than Qβ pilus detachment, MBP-Mat_Qβ_ can sufficiently detach F-pili, suggesting that the Mat protein can universally detach pili.

**Figure 3:**
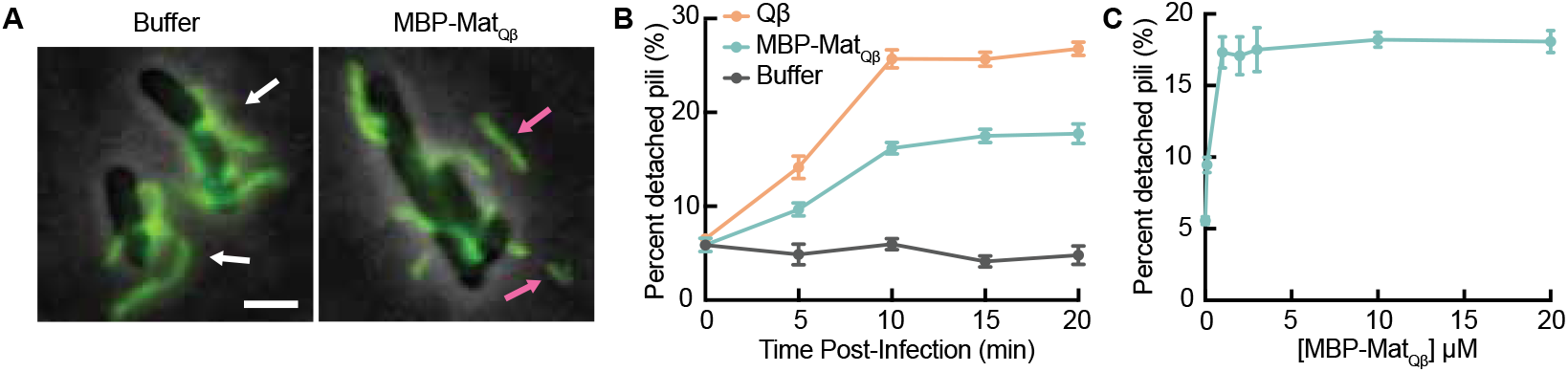
MBP-Mat_Qβ_ sufficiently induces F-pilus detachment. (A) Representative images of HfrH cells treated with MBP-Mat_Qβ_ or buffer. F-pilus were labeled by MS2-GFP. White arrows indicated cell-associated pili, pink arrows indicated detached pili in the background. (B) Percent of detached pili was quantified at different time points post-infection/treatment for Qβ, MBP-Mat_Qβ,_ and buffer. The number of total pili quantified at 0, 5, 10, 15, and 20 min post-infection for Qβ was 1306, 1197, 1266, 1131, and 1120, respectively; for MBP-Mat_Qβ_, 1199, 1237, 1214, 1293, and 1263, respectively; and for buffer, 1198, 1304, 1182, 1049, and 1104, respectively. Each time point was tested in triplicate. (C) Percent of detached pili was quantified after 10 min of treatment with different concentrations of MBP-Mat_Qβ_ (0, 0.1, 1, 2, 3, 10, and 20 μM). The number of total pili quantified for MBP-Mat_Qβ_ concentrations (0, 0.1, 1, 2, 3, 10, and 20 μM) was 1295, 1336, 1217, 1292, 1410, 1105, and 1175. Each concentration was tested in triplicate. Scale bar, 2 μm.

We then investigated the effect of MBP-Mat_Qβ_ concentration on F-pilus detachment. Cells were treated with increasing concentrations of MBP-Mat_Qβ_ for 10 min at which time the pilus detachment plateaus (**Fig. 3B**). The percentage of detached pili increased from a baseline of ∼5% without treatment to a peak of ∼17% at concentrations of 1 µM and above (**Fig. 3C**). Taken together with the results of Mat_200_ of AP205, we conclude that the phage Mat alone or the pilus binding domain of Mat is sufficient for pilus detachment.

### Mat_200_-Fluospheres can efficiently detach T4P

As shown in **Fig. 2D**, Mat_200_ can sufficiently detach T4P; however, the overall efficiency of this process is lower than that of the intact AP205 phage. We hypothesized that this difference in efficiency could be due to the structure and rigidity of the AP205 capsid, which can add extra load to break the pilus initiated by Mat while entering the cell. To test this, we conjugated Mat_200_ with a Fluosphere, which mimics the size and shape of the AP205 capsid. Specifically, we created a new protein construct, Mat_200_-AviTag (**Fig. 4A**). The biotinylated Mat_200_-AviTag was then conjugated to neutravidin-coated Fluospheres, generating a complex that mimics the size, shape, and binding of AP205 (**Fig. 4B**).

**Figure 4:**
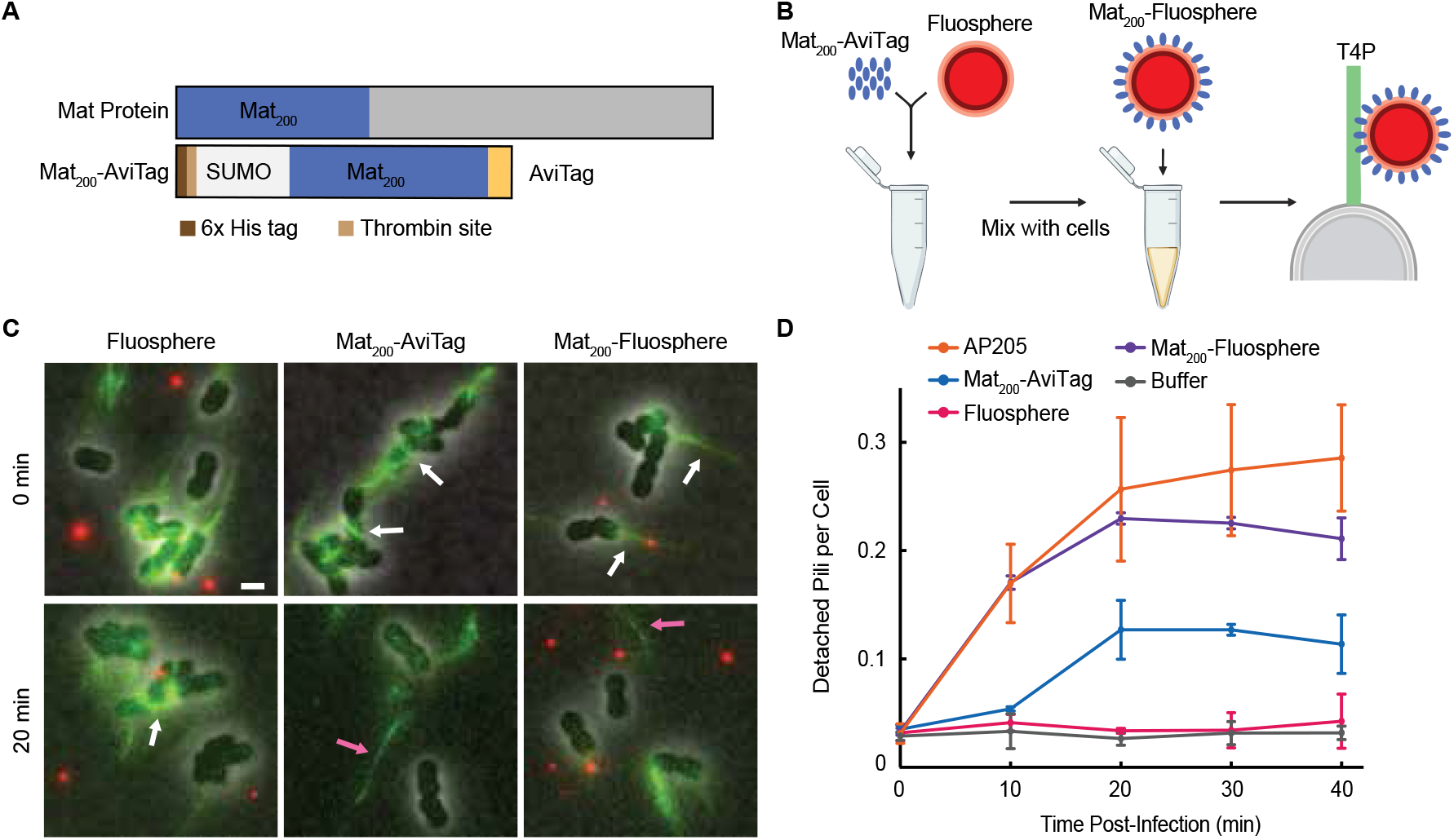
Mat_200_-Fluospheres can bind and detach T4P effectively. (A) Schematic representation of the full-length Mat protein versus the truncated Mat_200_-AviTag. This diagram illustrates the comparative sizes and domains of the two protein constructs. (B) Experimental schematic for assessing Mat_200_-Fluosphere generation and T4P interaction. The schematic is drawn to scale. Created in BioRender. Frese, A. (2026) https://BioRender.com/i66aujp (C) Representative micrographs illustrating the effects of various treatments on *A. higginsii* T4P. Images compare cells treated with Fluosphere, Mat_200_-AviTag, or Mat_200_-Fluosphere at 0 min and 20 min post-treatment. Fluospheres are visible as red puncta, while T4P are labeled with Mat_200_-sfGFP (green). White arrows indicate cell-associated pili, and pink arrows indicate detached pili. (D) Quantification of detached pili per cell at 0, 10, 20, 30, and 40 min post-infection for Mat_200_-AviTag, Fluosphere, and Mat_200_-Fluosphere. The total number of detached pili per cell at 0, 10, 20, 30, and 40 min post-treatment were for Fluosphere, 31/910, 60/1511, 32/986, 35/1083, and 39/1000, respectively; for Mat_200_-AviTag, 73/1899, 59/872, 171/1302, 60/489, and 103/663, respectively; for Mat_200_-Fluosphere, 73/1263, 249/1452, 237/1015, 205/927, and 240/948, respectively. The data for AP205 and buffer treated cells were taken from **Fig. 1B**. The total number of cells and detached pili were calculated across multiple experiments which were each performed in at least triplicate. Scale bar, 2 μm.

To test the pilus detachment, we treated *A. higginsii* cells with Mat_200_-AviTag, Fluospheres, or Mat_200_-Fluospheres (**Fig. 4C**). Treatment with Mat_200_-AviTag caused pilus detachment, reaching a plateau of ∼0.12 detached pili per cell after 20 min, a level similar to that of Mat_200_ and Mat_200_-sfGFP in **Fig. 2D** (**Fig. 4D**). This confirmed that the AviTag tag did not disrupt the ability of Mat_200_ to bind and detach T4P. Cells treated with Fluospheres alone showed a baseline level of ∼0.035 detached pili per cell, likely due to nonspecific interactions causing pilus shearing (**Fig. 4D**). When cells were treated with the Mat_200_-Fluosphere, the number of detached pili significantly increased, peaking at ∼0.23 detached pili per cell after 20 min, similar to the level by AP205 (**Fig. 4D**). This demonstrates that while Mat_200_ can detach pili sufficiently, it lacks efficiency. When Mat_200_ is bound to a large, rigid structure, this efficiency is restored to that of intact AP205 particles, indicating that Mat_200_ requires this structure to be fully effective.

## Discussion

In this study, we investigated the mechanism of pilus detachment by ssRNA phages using two distinct host-phage systems: AP205 of *A. higginsii* and Qβ of *E. coli*. We found that the ssRNA phage AP205 triggers a marked increase in pilus detachment, consistent with previous findings for MS2/Qβ and PP7^12,13^. The dose-dependent nature of pilus detachment, as shown by the increase in detachment with higher MOIs of AP205, suggests that it is a direct result of the phage infection process. This collective evidence from multiple host-ssRNA phage systems, including that of MS2/Qβ in *E. coli*, PP7 in *P. aeruginosa*, and AP205 in *A. higginsii*, strongly suggests that pilus detachment is a universally conserved mechanism essential for viral RNA entry.

Our findings identify Mat as the key viral component responsible for pilus detachment owing to its function of pilus binding (**Fig. 5**). We demonstrated that Mat_200_ sufficiently induces pilus detachment of *A. higginsii* T4P. This ability of a single viral protein to initiate a critical step of infection highlights the pivotal role of Mat in the vRNA delivery process. We also found that Mat_200_ can induce pilus detachment even when fused to a protein cargo like a sfGFP or an AviTag. This shows that Mat’s ability to bind and detach pili is retained even when it’s part of a larger complex, mimicking its natural state within the phage virion, where it’s bound to the viral RNA. The fact that the fusion proteins still caused significant detachment compared to buffer-treated samples confirms that the activity of Mat_200_ is not inhibited by the presence of a cargo.

**Figure 5:**
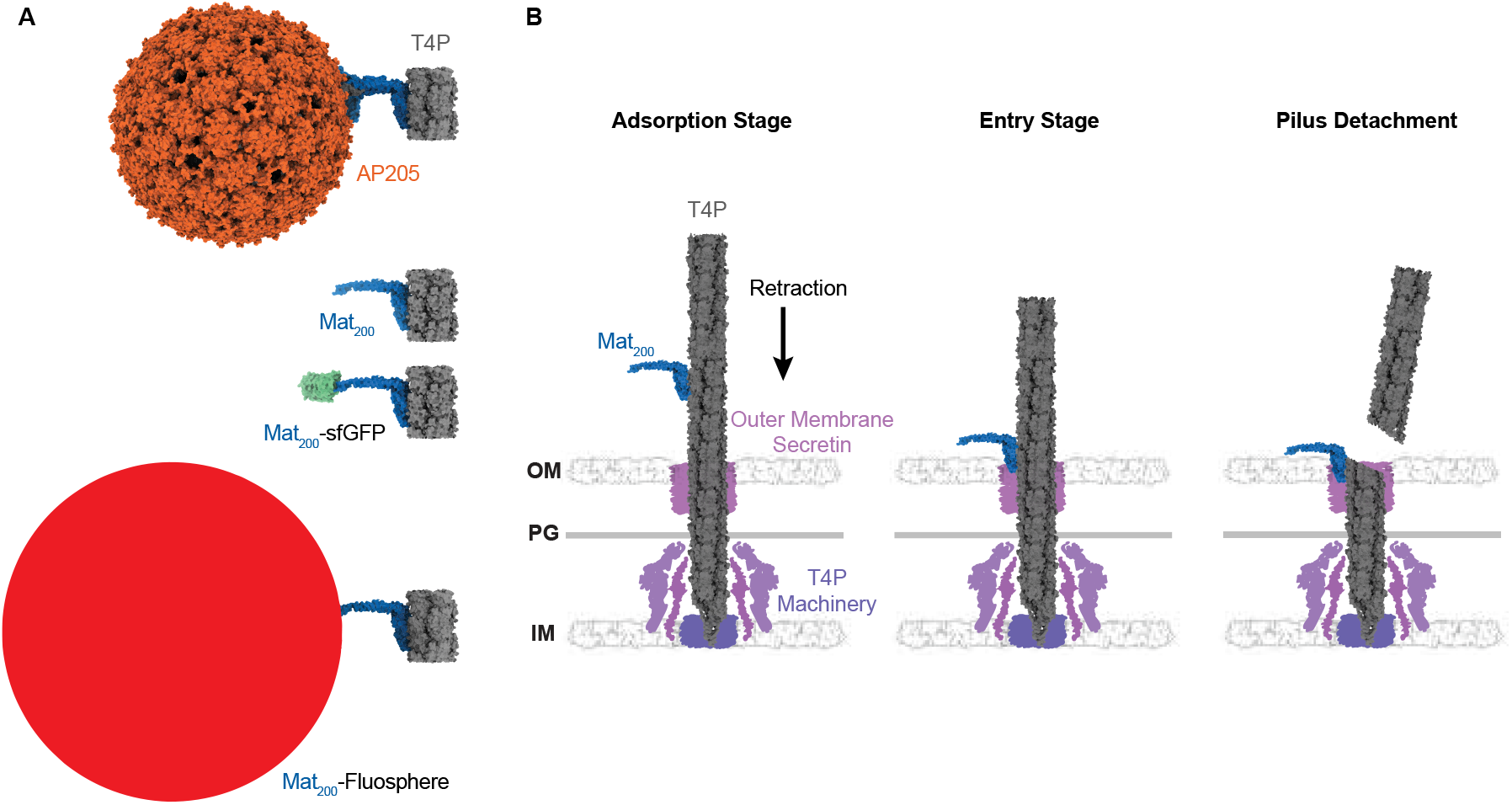
Model of T4P detachment as a result of engineered AP205 proteins. (A) Comparing the sizes of AP205, Mat_200_, Mat_200_-sfGFP, and Mat_200_-Fluosphere bound to T4P of *A. higginsii* (grey). Models are shown to scale. The composite models were created based on previously published PDB entries for AP205 (8TOC), Mat_200_ (8TVA), sfGFP (2B3P), and *A. higginsii* T4P (8TOB). (B) A schematic model that depicts the requirements for T4P detachment by Mat_200_. During the adsorption stage, Mat_200_ binds to a T4P. Retraction of a T4P brings Mat_200_ into proximity of the outer membrane secretin, leading to the entry stage. Once Mat_200_ enters the outer membrane secretin, the T4P detaches. Models are shown to scale.

Despite this finding, the efficiency of pilus detachment for either Mat_200_ in AP205 or Mat in Qβ was significantly less efficient than that of the entire ssRNA phage virion. This led us to hypothesize that the full efficiency of pilus detachment is dependent on the structural integrity of the phage virion. We created a Mat_200_-Fluosphere construct that mimic the size and rigidity of the AP205 virion to test our hypothesis. Critically, these constructs restored detachment efficiency to a level similar to that of the intact AP205 phage. This finding supports a model where the physical stress exerted on the pilus, caused by its retraction of a large, rigid, and bound structure, is the primary driver of detachment. The smaller, soluble Mat_200_ protein can initiate the detachment process but lacks the necessary physical properties to consistently overwhelm the pilus retraction machinery. When the Mat protein is part of the larger AP205 virion or a similarly sized Fluosphere, the forces generated during pilus retraction are sufficient to cause the pilus to shear or break at the cell surface, allowing for vRNA delivery.

## Methods

### Bacterial strains and plasmids

Bacterial strains, plasmids, and oligonucleotides used in this study are listed in **Table S1**. *A. higginsii* cells were grown in LB medium in a 14 mL Falcon round-bottom tube at 30°C with shaking (225 rpm) unless indicated otherwise. All *E. coli* strains were grown in LB medium containing related antibiotics, in baffled flasks at 37°C with shaking (265 rpm) unless other conditions indicated.

### AP205 Purification

A single colony of *A. higginsii* cells was grown in 25 mL of LB overnight at 30°C with shaking at 225 rpm in a 250 mL round-bottom flask. The overnight culture was diluted 1:100 in 1 L of fresh LB in a 4 L round-bottom flask and grown at 30°C with shaking (225 rpm) until the OD_600_∼0.1. 300 µL of AP205 phage lysate (∼10^10^ PFU/mL) was added to the culture supplemented with 2 mM CaCl_2_. After the phage is added, the culture is allowed to sit at room temperature for 30 min without shaking before the culture is grown. The culture was grown overday (∼7-8 hrs) before the supernatant was harvested. The cells were split into two 1 L centrifuge bottles and were spun down at 13,000 × g for 30 min at 4°C in an F9-6x1000 fixed-angle rotor to separate the supernatant lysate and cell pellet. The supernatant lysate was passed through a 0.22 μm filter to remove excess cellular debris. Next, ∼950 mL of phage lysate was precipitated with ammonium sulfate using 280 g per liter of phage lysate. ∼20 g of ammonium sulfate was added at a time, with 20 min between each addition, with stirring to allow the solution to dissolve fully. The precipitated phages were harvested via centrifugation at 14,330 × g for 1 hr at 4°C in a 1 L centrifuge bottle in an F9-6x1000 fixed-angle rotor. Once the supernatant was discarded, the pellet was resuspended in AP205 buffer (150 mM NaCl, 50 mM Tris-HCl (pH 8), 5 mM EDTA) and dialyzed against 2 L of AP205 buffer a total of three times for at least 3 hrs each time using a 20 kDa MWCO dialysis cassette. The phage was then subjected to CsCl isopycnic ultracentrifugation with the addition of CsCl in a 13 mL quick-seal centrifuge tube to a final density of 1.38 g/mL, then spun at 45,000 rpm for 24 hrs using a Ti 70 rotor. Next, the bands of phage that formed were removed using an 18-gauge needle and dialyzed against 1 M NaCl for 5 hrs using a new 20 kDa MWCO dialysis cassette. This was followed by dialyzing against 1 L of AP205 buffer a total of three times for 8 hrs, overnight, and 8 hrs.

### Pilus Detachment Assay and Fluorescence Imaging

An overnight culture containing a single colony of *A. higginsii* cells was grown in 3 mL LB inside of a 14 mL Falcon round-bottom tube at 30°C with shaking (225 rpm) for ∼16 hrs. The following morning, an overday culture was grown by diluting the overnight culture 1:100 into 3 mL of fresh LB in a 14 mL Falcon round-bottom tube until the cells reached an OD_600_ ∼ 0.4 (∼3.5-4 hrs). 40 µL of cells were aliquoted into microcentrifuge tubes and treated with 5 µL of the respective treatment agent (AP205 phage at MOI = 5, Mat_200,_ Mat_200_-sfGFP, Mat_200_-AviTag, Mat_200_-Fluosphere, or buffer control). Samples were incubated in a 30°C water bath without shaking for defined time intervals (0, 2, 5, 10, 20, 30, or 40 min). Immediately following treatment, 5 µL of Mat_200_-sfGFP (15 μM) was added to label the pili for 2 min at room temperature in dark conditions. A small drop (∼1 µL) of the labeled sample was added onto a 1.5% PBS agarose pad (∼1 mm thick) on top of a small coverslip (18 × 18 mm), which was topped with a large coverslip (24 × 50 mm). Imaging was performed at room temperature using a Nikon Eclipse Ti2 inverted epifluorescence microscope equipped with a 100x objective lens (Plan Apochromat, numerical aperture 1.45, oil immersion) and a cooled EMCCD camera with a mask (Princeton Instruments). Z-stacks were captured at 300 nm intervals. Exposure times were set as follows: phase contrast (100 ms), GFP channel (10 ms) to visualize labeled pili (Chroma 96362), and mCherry/TxRed channel (40 ms) to detect Fluospheres (Chroma 96365). Images were analyzed using a combination of NIS-Elements software (Nikon), OmniPose for cell segmentation and counting, ImageJ FIJI, and the MicrobeJ plugin for detached pili identification and counting^17–19^.

For the pilus detachment experiments involving Mat_200_, Mat_200_-sfGFP, Mat_200_-AviTag, or Mat_200_-Fluospheres, the methods were kept the same with a few exceptions. Cell growth conditions remained the same; however, 40 µL of cells were treated with 5 µL of either Mat_200_, Mat_200_-sfGFP, Mat_200_-AviTag, or Mat_200_-Fluospheres before incubation at 30°C for varying times. Pilus labeling and imaging conditions were kept the same.

### Mat_200_-sfGFP, Mat_200_, and Mat_200_-AviTag Expression and Purification

The expression and purification of Mat_200_-sfGFP and Mat_200_ were performed based on previously established methods^10^. Briefly, *E. coli* BL21 DE3 cells containing either the pET28-His-SUMO-Mat_200_-sfGFP or the pET28-His-SUMO-Mat_200_ vector were grown overnight at 37°C with shaking (200 rpm) with 25 mL LB and 50 µg/mL of kanamycin in a 250 mL baffled flask. The overnight culture was added to 1 L of fresh LB with kanamycin in a 4 L baffled flask for expression assays and was grown at 37°C with shaking (200 rpm) until the OD_600_ reached 0.4-0.6. The cells were chilled on ice and transferred to 16°C with shaking (200 rpm) and induced with 0.5 mM IPTG overnight (∼16 hrs). The cells were harvested using centrifugation at 4,000 × g for 30 min at 4°C in a 1 L centrifuge bottle. Once the supernatant was removed, the pellet was resuspended in 50 mL of lysis buffer (50 mM Tris-HCl (pH 8), 200 mM NaCl) containing a Pierce™ Protease Inhibitor Tablet (ThermoFisher, A32965). The cells were lysed using a microfluidizer at 25,000 psi, and the supernatant was collected using centrifugation at 14,000 × g for 30 min at 4°C in a 50 mL round-bottom centrifuge tube. The collected supernatant was injected into a Nickel affinity column, which was washed with washing buffer (20 mM Tris-HCl (pH 8), 200 mM NaCl, 5 mM Imidazole), and His-SUMO-Mat_200_-sfGFP was eluted with elution buffer (20 mM Tris-HCl (pH 8), 200 mM NaCl, 300 mM Imidazole). The eluted protein was concentrated using a 10K MWCO protein concentrator and spun at 4,700 × g for 60 min at 4°C. 500 µL of concentrated protein was loaded into a Superdex 200 size-exclusion column using an equilibration buffer (50 mM Tris-HCl (pH 8), 200 mM NaCl) for elution. The fraction that corresponds to the His-SUMO-Mat_200_-sfGFP or His-SUMO-Mat_200_ protein was collected and confirmed by an SDS-PAGE gel. A commercial SUMO tag protein (Proteogenix, ARO-P12692) was used as a negative control in pili detachment assays to assess the potential influence of the purification tag.

For the production of biotinylated protein, a similar protocol was followed as above for Mat_200_-sfGFP expression and purification, except with several modifications. To begin, the pET28-His-SUMO-Mat_200_-AviTag plasmid was generated using the pET28-His-SUMO-Mat_200_-sfGFP plasmid. Gibson assembly was used to replace the sfGFP with an AviTag (**Table S2**). The pET28-His-SUMO-Mat_200_-AviTag vector was co-transformed with a BirA-expressing plasmid, pBAD-birA, into *E. coli* BL21 DE3 cells. The cells were grown in LB supplemented with 50 µg/mL kanamycin and 10 µg/mL chloramphenicol. At an OD_600_ ∼ 0.4-0.6, the culture was supplemented with 50 µM D-biotin and protein expression was induced with 0.5 mM IPTG overnight at 16°C with shaking (200 rpm). Subsequent cell harvesting, lysis, and protein purification steps were performed as described above.

### Mat_200_-Fluosphere Generation, and Fluorescent Imaging

Purified Mat_200_-AviTag was stored at 4°C at a concentration of ∼4 × 10^15^ particles/mL. Neutravidin-coated Fluospheres (Thermo Fisher, F8770) were stored at 4°C at a concentration of 4 × 10^12^ particles/mL. The Fluospheres are 35 ± 5 nm in diameter and have red fluorescence (580/605 nm). In a microcentrifuge tube, 25 μL of the diluted Fluospheres at 4 × 10^10^ particles/mL, 25 μL of diluted biotinylated Mat_200_-AviTag at 4 × 10^12^ particles/mL, and 50 μL of PBS were mixed. The mixture was allowed to incubate for 1 hr at room temperature in dark conditions. Any unbound Mat_200_-AviTag was washed by adding the sample to a 100 kDa MWCO spin column and centrifuging at 4,000 × g for 15 min at room temperature. The flow-through contains the unbound Mat_200_-AviTag, while the Mat_200_-Fluospheres are contained above the filter.

### MBP-Mat_Qβ_ expression and purification

The expression and purification of MBP-Mat_Qβ_ were performed based on previously reported^16^. Briefly, pET28-MBP-Qβ was transformed into the BL21 strain, which originally contains the pZA32-MurAA plasmid to prevent Qβ-caused cell lysis. Cells were grown at 37°C with 265 rpm shaking until OD_600_ reached 0.6, then chilled in an ice bath for 1 hr before induction with 1 mM IPTG and expression at 16°C, 180 rpm for 18 hrs. Cells were harvested by centrifugation at 12,000 × g for 15 min at 4°C and resuspended in lysis buffer [20 mM Tris-HCl (pH 8), 150 mM NaCl with 1% glycerol, 1 mM EDTA, 20 μg/mL DNase, 10 μg/mL RNaseA, and Pierce™ Protease Inhibitor Tablet (Thermo Fisher, A32965)]. Cells were disrupted by passage through an SLM-Aminco French pressure cell (Spectronic Instruments) at 16,000 psi, and lysate was clarified by centrifugation at 12,000 × g for 15 min at 4°C, and then filtered through a 0.45 μm filter before loading onto an amylose resin column (NEB cat #E8021S). Protein was eluted with 10mM maltose in elution buffer (20 mM Tris-HCl (pH 8), 150 mM NaCl, 1% glycerol).

### Labeling of F-pilus

The visualization of F-pilus was performed by incubating with MS2-GFP. The expression and purification of MS2-GFP was performed as previously reported^13^. Briefly, 10 mL of LZ2619 overnight culture was grown in a 100 mL flask was diluted 1:100 into 1 L fresh LB. Then, 800 mL of the LZ2619 overday culture was evenly separated into two 4 L flasks, and the remaining 200 mL of the overday culture was separated into one 2 L flask. These cultures were grown at 37°C with shaking (265 rpm) for ∼ 2.5 hrs until the OD_600_ ∼ 0.6, before the MS2 lysate was then added at an MOI = 5. The flasks were then left on the bench top at room temperature for 10 min, allowing phages to adsorb to cells. After that, a final concentration of 1 mM IPTG was added, and the culture was allowed to continue to grow at the previous conditions. After 6 hrs, the supernatant was harvested by spinning the culture down at 8,000 × g for 30 min at 4°C. Purification of the phages was achieved using ammonium sulfate precipitation and CsCl gradient centrifugation. In brief, 280 g/L of ammonium sulfate was added to the lysates in small increments, followed by chilling at 4°C for 4 hrs. The precipitates were sedimented by centrifugation at 12,000 × g for 1 hr and resuspended in MS2 buffer (150 mM NaCl, 5 mM EDTA, and 50 mM Tris-HCl (pH 7.5)). The suspensions were dialyzed in 10 kDa MWCO cassettes three times at 2 hr intervals with MS2 buffer at 4°C, followed by treatment with DNase at 10 U/mL for 1 hr at 4 °C. A clearing spin (8,000 x g for 1 hr) was performed, after which the phage preparations were mixed with 0.55 g CsCl per gram of phage solution to produce a final density of 1.38 g/mL. The suspensions were centrifuged at 45,000 rpm for 24 hrs at 4°C. The bands were collected and dialyzed against 2 L of MS2 buffer (with 1M NaCl) for 6 hrs, then replaced with fresh 2 L of MS2 buffer (with 250 mM NaCl) for dialyzing overnight.

### F-pilus detachment assay

Overnight cultures of piliated cells were diluted 1:100 in LB containing antibiotics and 4 mM CaCl_2_. The cells were grown to an OD_600_ of 0.4, then purified Qβ or MBP-Qβ was added to the culture. Infections were performed at 37 °C without shaking to minimize pilus agitation. Samples were removed at 0, 5, 10, 15, and 20 min using cut pipette tips and dispensed into prechilled tubes and allowed to equilibrate on ice. A volume of MS2-GFP equating to an MOI of 20 was added to the samples and left on ice for 20 min to allow phage adsorption to the pili. F-pilus retraction is significantly inhibited at low temperature, and the addition of MS2-GFP does not influence the detachment of pili^13^.

Samples were prepared for microscopy by spotting 1 μL of sample onto a large coverslip (No. 1, 24 × 50 mm; Thermo Fisher Scientific) and gently overlaying a small, 1-mm-thick 1.5% agarose pad dissolved in PBS on the sample before placing it under the microscope. Z stacks of 300 nm were used in the GFP channel to precisely measure pili. Cells and pili were imaged on multiple stage positions (∼10-20 per sample) in phase (100 ms exposure to detect cells) and GFP (100 ms to detect pili) channels. The images were analyzed using NIS-Elements software (Nikon) and Fiji for cell and pili counts and length measurements.

## Acknowledgments

We would like to thank the Junjie Zhang laboratory for gifting strains along with Wen Xiao and Zhongliang Xing for assisting in protein purification. We are grateful for Ry Young and the members of the Zeng, Zhang, Hays Rye, Jason Gill, and Jolene Ramsey laboratories for discussions. This work was supported by the National Institutes of Health grant R01GM141659.

## References

1. Castanheira, M., Mendes, R.E. & Gales, A. C. Global Epidemiology and Mechanisms of Resistance of Acinetobacter baumannii-calcoaceticus Complex. Clin. Infect. Dis. Off. Publ. Infect. Dis. Soc. Am. 76, S166–S178 (2023).

2. Iwashkiw, J. A. et al. Identification of a General O-linked Protein Glycosylation System in Acinetobacter baumannii and Its Role in Virulence and Biofilm Formation. PLOS Pathog. 8, e1002758 (2012).

3. Antunes, L. C. S., Visca, P. & Towner, K. J. Acinetobacter baumannii: evolution of a global pathogen. Pathog. Dis. 71, 292–301 (2014).

4. CDC. 2019 Antibiotic Resistance Threats Report. Antimicrobial Resistance https://www.cdc.gov/antimicrobial-resistance/data-research/threats/index.html (2025).

5. Wall, D. & Kaiser, D. Type IV pili and cell motility. Mol. Microbiol. 32, 01–10 (1999).

6. Hobbs, M. & Mattick, J. S. Common components in the assembly of type 4 fimbriae, DNA transfer systems, filamentous phage and protein-secretion apparatus: a general system for the formation of surface-associated protein complexes. Mol. Microbiol. 10, 233–243 (1993).

7. Jacobsen, T., Bardiaux, B., Francetic, O., Izadi-Pruneyre, N. & Nilges, M. Structure and function of minor pilins of type IV pili. Med. Microbiol. Immunol. (Berl.) 209, 301–308 (2020).

8. Ellison, C. K., Whitfield, G.B. & Brun, Y. V. Type IV Pili: dynamic bacterial nanomachines. FEMS Microbiol. Rev. 46, fuab053 (2022).

9. Burrows, L. L. Pseudomonas aeruginosa Twitching Motility: Type IV Pili in Action. Annu. Rev. Microbiol. 66, 493–520 (2012).

10. Meng, R. et al. Structural basis of Acinetobacter type IV pili targeting by an RNA virus. Nat. Commun. 15, 2746 (2024).

11. Fernandez-Martinez, D. et al. Cryo-EM structures of type IV pili complexed with nanobodies reveal immune escape mechanisms. Nat. Commun. 15, 2414 (2024).

12. Thongchol, J. et al. Removal of Pseudomonas type IV pili by a small RNA virus. Science 384, eadl0635 (2024).

13. Harb, L. et al. ssRNA phage penetration triggers detachment of the F-pilus. Proc. Natl. Acad. Sci. 117, 25751–25758 (2020).

14. Klovins, J., Overbeek, G. P., van den Worm, S. H. E., Ackermann, H.-W. & van Duin, J. Nucleotide sequence of a ssRNA phage from Acinetobacter: kinship to coliphages. J. Gen. Virol. 83, 1523–1533 (2002).

15. Nemec, A. et al. Proposal for Acinetobacter higginsii sp. nov. to accommodate organisms of human clinical origin previously classified as Acinetobacter genomic species 16. Int. J. Syst. Evol. Microbiol. 73, 006114 (2023).

16. Reed, C. A., Langlais, C., Kuznetsov, V. & Young, R. Inhibitory mechanism of the Qβ lysis protein A2. Mol. Microbiol. 86, 836–844 (2012).

17. Cutler, K. J. et al. Omnipose: a high-precision morphology-independent solution for bacterial cell segmentation. Nat. Methods 19, 1438–1448 (2022).

18. Schindelin, J. et al. Fiji: an open-source platform for biological-image analysis. Nat. Methods 9, 676–682 (2012).

19. Ducret, A., Quardokus, E.M. & sBrun, Y. V. MicrobeJ, a tool for high throughput bacterial cell detection and quantitative analysis. Nat. Microbiol. 1, 16077 (2016).

